# Task errors drive memories that improve sensorimotor adaptation

**DOI:** 10.1101/538348

**Authors:** Li-Ann Leow, Welber Marinovic, Aymar de Rugy, Timothy J Carroll

**Author notes:** Corresponding author: Li-Ann Leow. Corresponding author. **Significance Statement**. Latent motor memories formed during sensorimotor adaptation manifest as improved re-learning when sensorimotor perturbations are re-encountered (i.e., savings). Conflicting theories suggest that savings is underpinned by different mechanisms: including a memory of successful actions; a memory of errors; or an aiming strategy to correct task errors. To elucidate this issue, we examined the necessary conditions for savings. We show that learning to correct task errors is sufficient for savings, even when tested in the absence of task errors. In contrast, a history of sensory prediction errors is neither sufficient nor obligatory for savings. Finally, we show that latent sensorimotor memories driven by task errors comprise at least two distinct components: a time-consuming, flexible component, and a rapidly expressible, inflexible component.

## Abstract

Traditional views of sensorimotor adaptation, or adaptation of movements to perturbed sensory feedback, emphasize the role of automatic, implicit correction of sensory prediction errors (differences between predicted and actual sensory outcomes). However, latent memories formed during sensorimotor adaptation, manifest as improved learning (i.e., savings), have recently been attributed to strategic corrections of task errors (failures to achieve task goals). To dissociate contributions of task errors and sensory prediction errors to latent sensorimotor memories, we perturbed target locations to remove or enforce task errors during learning and/or test. We show that prior learning to correct task errors alone, even in the absence of perturbation-induced sensory prediction errors, was sufficient for savings, even when tested in the absence of task errors. In contrast, a history of sensory prediction errors was neither sufficient nor obligatory for savings. Limiting movement preparation time further revealed that the latent memories that are driven by learning to correct task errors take at least two forms: a time-consuming but flexible component, and a rapidly expressible, inflexible component. The results provide strong support for the idea that failure to successfully achieve movement goals is the primary driver of motor memories that manifest as savings. In contrast to previous suggestions, such persistent memories are not exclusively mediated by time-consuming strategic processes, but also comprise a rapidly expressible but inflexible component. The distinct characteristics of these putative processes that contribute to persistent motor memories suggest dissociable underlying mechanisms, and imply that identification of the neural basis for adaptation and savings will require methods that allow such dissociations.

## Introduction

Much of what we know about motor learning emerged from the paradigm of sensorimotor adaptation, where a systematic perturbation is applied to the visual representation of movement (von Helmholtz and Southall, 1924; Cunningham, 1989), or to limb dynamics (Dietz et al., 1994; Shadmehr and Mussa-Ivaldi, 1994). This changes the sensory consequences of motor commands, prompting adaptive motor responses to restore effective movement. Adaptation is essential to sustain movement success, due to ubiquitous changes within our bodies and the environment. However, the extent to which principles of sensorimotor adaptation apply to learning of motor skills remains unclear (Krakauer and Carmichael, 2017).

A key piece of evidence addressing the question of how sensorimotor adaptation relates to other forms of motor learning is the extent to which adaptation generates long-lasting memories. Sensorimotor memory is often operationalised as improved adaptation when re-encountering a similar perturbation. This phenomenon, commonly termed savings, implies latent memories (Ebbinghaus, 1964), because the benefit of previous learning persists for months even after behaviour has returned to the naïve state (Zarahn et al., 2008; Landi et al., 2011; Maeda et al., 2018). A latent form of sensorimotor memory is both obligatory for success given the non-stationarity of our environment, and reminiscent of learnt motor skills, which can be flexibly expressed according to context.

The mechanisms that underlie savings are unresolved. One idea suggests that savings occurs because specific actions become associated with success during learning, and these actions are recalled upon perturbation-induced failure (Huang et al., 2011). Another idea suggests a role of “meta-learning”, where the structure of an experienced perturbation is encoded to assist subsequent adaptation (Braun et al., 2009). Yet another influential theory suggests that a systematic sequence of perturbation-induced errors generates a memory of errors, increasing sensitivity to those errors and the gain of error correction when reencountering similar errors (Gonzalez Castro et al., 2014; Herzfeld et al., 2014). In contrast, recent work provides strong evidence that savings is dominated by strategic selection of actions that restore success (Haith et al., 2015; Morehead et al., 2015; de Brouwer et al., 2017; Avraham et al., 2019). But is strategy use the sole contributor to latent sensorimotor memories, or is there some component of latent sensorimotor memories that are less accessible to conscious control? If there are indeed multiple components to sensorimotor memories, what are the necessary conditions for their encoding and expression? Sensorimotor perturbations, by definition, evoke sensory prediction errors (SPEs, or, discrepancies between predicted sensory outcomes of movements and actual sensory outcomes of movements). Perturbations also often evoke reward prediction errors (RPEs, or discrepancies between predicted and actual reward). In a goal-directed task, reward prediction errors take the form of task error, or failure to achieve an internally determined task goal. Previous conceptionalizations delineate the neural mechanisms of SPEs and RPEs, where the cerebellum uses SPEs to improve action execution, while the basal ganglia uses RPEs to improve action selection (Taylor and Ivry, 2014). However, recent evidence not only shows reciprocal connections between the two systems (Bostan and Strick, 2018), but also that the function of these systems is not completely dissociable (e.g., both SPEs and RPEs modulate cerebellar responses (Heffley et al., 2018; Kostadinov et al., 2019)). In sensorimotor learning, these errors can interact to influence adaptation (e.g., Leow et al., 2018; Pidoux et al., 2018; Kim et al., 2019). Could SPEs and RPEs jointly contribute to latent sensorimotor memories? To answer this question, we examined the contibution of task errors and SPEs to the encoding and expression of sensorimotor memories. We show that a history of adaptation to task errors, and not a history of adaptation to SPEs, is necessary to encode memories that subsequently improve adaptation. Further, we show that this memory consists of at least two components: a time-consuming but flexible component that readily accommodates different task demands, and a rapidly-expressible, inflexible component that fails to accommodate new task demands.

## Methods and Materials

### Participants

There were a total of 132 participants (75 female, age range 17-34 years, mean age 20.6). All participants were naïve to visuomotor rotation and force-field adaptation tasks, and were naïve to the aims of the study. Participants received course credit or monetary reimbursement upon study completion. The study was approved by the Human Research Ethics Committee at The University of Queensland. All participants provided written informed consent. This study conforms with the Declaration of Helsinki.

### Apparatus

Participants completed the task using a vBOT planar robotic manipulandum, which has a low-mass, two-link carbon fibre arm and measures position with optical encoders sampled at 1,000 Hz (Howard et al., 2009). Participants were seated on a height-adjustable chair at their ideal height for viewing the screen for the duration of the experiment. Visual feedback was presented on a horizontal plane on a 27” LCD computer monitor (ASUS, VG278H, set at 60Hz refresh rate) mounted above the vBOT and projected to the participant via a mirror in a darkened room, preventing direct vision of her/his hand. The mirror allowed the visual feedback of the targets, the start circle, and hand cursor to be presented in the plane of movement, with a black background. The start was aligned approximately 10cm to the right of the participant’s mid-sagittal plane at approximately mid-sternum level. An air-sled was used to support the weight of participants’ right forearms, to reduce possible effects of fatigue.

### General Trial Structure

While grasping the robot arm, participants moved their hand-cursor (0.5cm radius red circle) from the central start circle (0.5cm radius white circle) to the targets (0.5cm radius yellow circles). Targets appeared in random order at one of eight locations (0°, 45°…. 315°) at a radius of 9 cm from a central start circle. At the start of each trial, the central start circle was displayed. If participants failed to move their hand-cursor to within 1cm of the start circle after 1 second, the robotic manipulandum moved the participant’s hand to the start circle (using a simulated 2 dimensional spring with the spring constant magnitude increasing linearly over time). A trial was initiated when the cursor remained within the home location at a speed below 0.1cm/s for 200ms. Across all experiments, we used a classical timed-response paradigm (e.g., e.g., Schouten and Bekker, 1967) to manipulate movement preparation time during the planar reaching task (Favilla and De Cecco, 1996). A sequence of three tones, spaced 500ms apart, was presented at a clearly audible volume via external speakers. Participants were instructed to time the onset of their movements with the onset of the third tone, which was more highly-pitched than the two previous, and then to slice through the target with their cursor. Movement initiation time was identified online as when hand speed exceeded 2cm/s. Targets appeared at 1000ms (Experiments 1-3) or 250ms (Experiment 4) minus a monitor display latency (27.6 ± 1.8ms), before the third tone. Thus, target direction information became available 972ms before the desired initiation time for Experiments 1-3 and 222ms before the desired initiation time for Experiment 4. When movements were initiated 50ms later than the third tone, the trial was aborted: the screen went black and the text “Too Late” was displayed on the feedback screen. When movements were initiated more than 100ms before the desired initiation time, the trial was aborted: the screen went black and a “Too Soon” error message was displayed. Thus, movements had to be initiated between 872 and 1022ms of target presentation in Experiments 1-3 and between 122ms and 272ms of target presentation in Experiment 4. We chose this movement preparation time for consistency with our previous work using the timed-response paradigm with visuomotor rotations (Leow et al., 2017). No visual feedback about movements was available when trials were aborted, and so such trials were repeated at the end of the cycle. We enforced long movement preparation times across Experiments 1-3 to prevent the possibility that the task error manipulation resulted in self-selection of different movement preparation times. Under these conditions, participants had ample opportunity (i.e. time for movement preparation) to use explicit strategies.

Across all conditions, cursor feedback was displayed after the hand had moved 4cm from the start to target (located 9cm away from the start). At this point, the direction of cursor velocity was measured to define target movements in some conditions as described below. During **StandardTaskError** conditions, the target remained stationary throughout the trial, such that whether or not participants hit the target was contingent on the participant’s reach direction (Figure 1A). During **NoTaskError** conditions, the target was shifted to align with the direction of cursor velocity, measured at 4cm into the movement. This is analogous to moving a basketball hoop towards the ball mid-flight; the ball always goes through the hoop regardless of the person’s actions. During **EnforcedTaskError** conditions, the target was shifted randomly by 20°–30° (counterclockwise in half of the trials, clockwise in half of the trials) relative to the cursor direction when the hand had moved 4 cm from the start. This is analogous to moving a basketball hoop away from the ball’s trajectory; participants can never get the ball through the hoop regardless of where they shoot. In Experiments 3 and 4, we imposed systematic task errors without any perturbation of the hand-cursor relationship: the target was moved during the movement by 30° relative to the original target position, always in the same direction (clockwise for half of all participants, counterclockwise for the other half of participants, counterbalanced): no rotation of the visual feedback of movement was imposed when this occurred.

**Figure 1.**
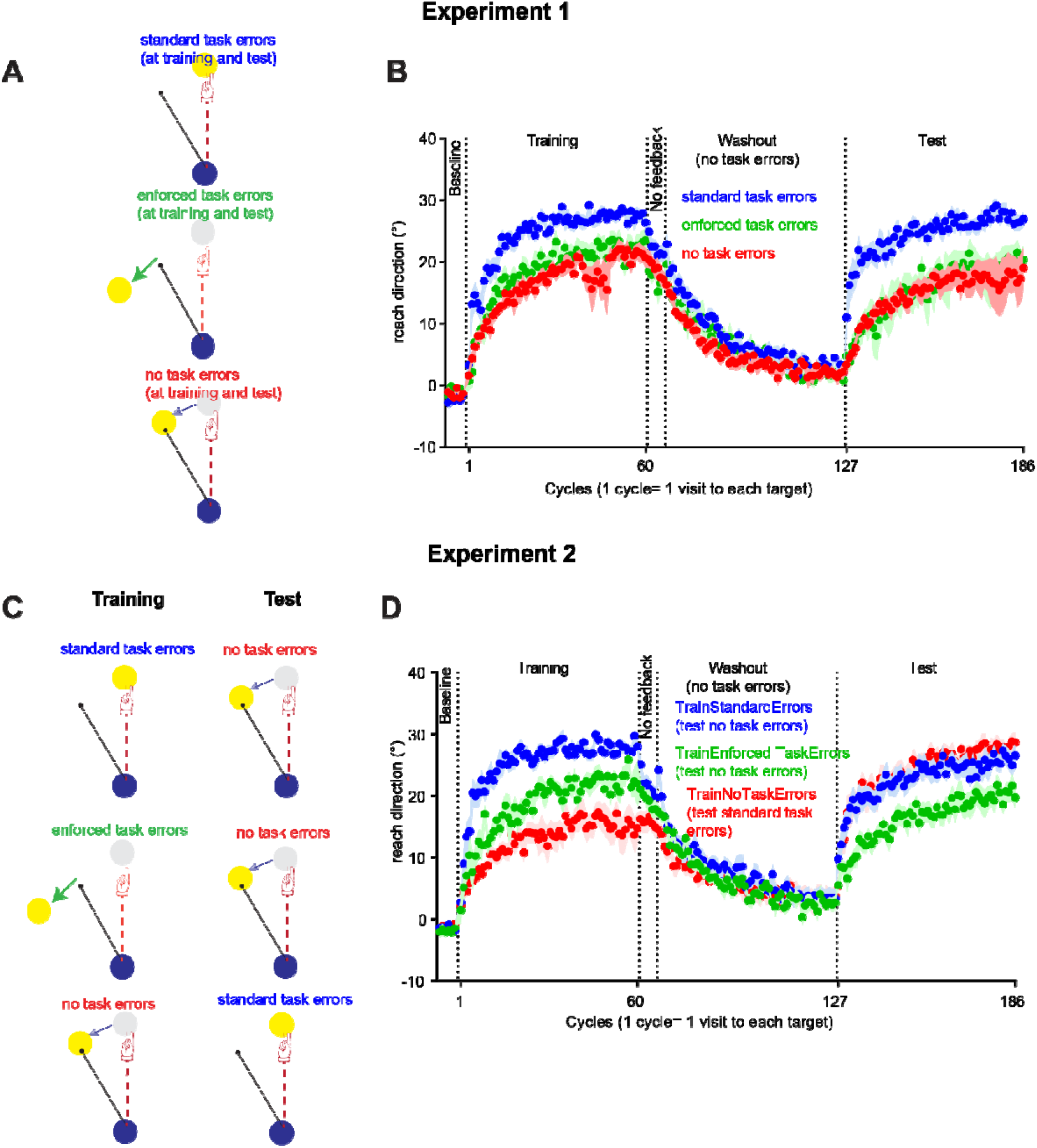
Task error manipulations in Experiment 1 (A) and Experiment 2 (C). All groups in Experiments 1 and 2 experienced the no-task error manipulation at washout. Cycle-averaged data for all experimental phases in Experiment 1 (B) and Experiment 2(D).

To familiarize participants with the equipment and the timed-response paradigm, all participants were first allowed a familiarization block comprising a maximum of 6 cycles. One cycle consisted of 1 trial to each of the 8 targets, and target order was random within each cycle. Participants were explicitly instructed to make shooting movements so that the cursor sliced through the targets, rather than to stop on the targets. Cursor feedback terminated as soon as the desired movement extent (the 9cm distance between the start and the target) was achieved. After familiarisation, all participants (regardless of assigned condition) were given the same task instruction, as follows. “Your task in this experiment is to hit the targets. The computer might disturb the cursor and/or the target, this is a normal part of the experiment, just try to hit the target as well as you can”. Participants then completed the following blocks. **Baseline** (6 cycles): no rotation of visual feedback. **Training** (60 cycles): For experiments 1 & 2, a 30° rotation of cursor feedback representing the hand position was imposed. Half of all participants encountered a clockwise 30°cursor rotation and half encountered a 30° counterclockwise cursor rotation. For experiments 3-4, no cursor rotation was applied during training, but a 30° rotation of target position relative to the original target position was applied mid-movement (Figure 4A). Half of all participants encountered a clockwise 30° target rotation and half encountered a 30° counterclockwise target rotation. **No feedback** (6 cycles): Upon leaving the start circle, no feedback about movements was available. Before this block, participants received explicit instructions about the rotation removal, as follows: “Any disturbance that the computer has applied is now gone, and the feedback of your movement will now be hidden as soon as it leaves the start circle, so please move straight to the target”. **Washout**: Cursor position feedback was restored, but the 30° rotation of cursor (or target) was removed. For Experiments 1 and 2, to prevent the experience of washout-related task errors, task errors were removed across all conditions (i.e., the target position shifted mid-movement to ensure that the cursor always hit the target). The length of the washout block was the same as the adaptation block (60 cycles). For Experiments 3 and 4, participants had no prior experience of the cursor rotation, only task errors, and they could volitionally reach straight to the target by the end of the no-feedback block, thus it was unnecessary to employ a long washout with the no-task-error manipulation to avoid exposure to errors related to abrupt removal of the perturbation: we thus provided 12 washout cycles without mid-movement target shifts. **Test** (60 cycles): the 30° rotation of cursor feedback was imposed (half of all participants encountered a clockwise 30° rotation and half encountered a 30° counterclockwise rotation). Between each block, there was a small delay to allow for experimental instructions and loading of the computer code for different experimental blocks.

### Data analysis

Movement onset time was taken as the time at which hand speed first exceeded 2 cm/s. Reach directions were quantified at 20 percent of the radial distance between the start and the target. Reaches with absolute initial direction errors greater than 60° with respect to the target (movements that were more than 60° to the left or the right of the target) were considered outliers, and were removed from analyses. Experiment1: StandardTaskErrors: 0.62%, NoTaskErrors 0.11%, EnforcedTaskErrors: 1.73%; Experiment2: TrainStandardTaskErrors: 0.29%, TrainEnforcedTaskErrors: 0.62%; TrainNoTaskErrors: 0.17%; Experiment 3: Same: 0.30%, Different: 0.20%. Experiment 4:ShortDifferent: 3.82%; ShortSame: 4.31%, ShortNaive:4.59%). Excluding these trials did not have any qualitative impact on the results. Trials were averaged in cycles of eight (one trial for each target angle) for conversion to percent adaptation (see below). For graphing purposes, reach directions for participants who experienced counterclockwise rotations (30°) were sign-transformed and pooled with data for participants who experienced counterclockwise (30°) rotations: values closer to 30° indicate more complete adaptation.

For all blocks except the test block, we estimated adaptation performance as percent adaptation, which quantifies reach directions relative to the ideal reach direction (as shown in Hadjiosif and Smith, 2013).

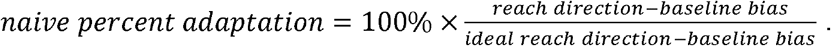

For the test block, we were interested in savings, which is improved learning at test compared to naïve. Even with exposure to the same number of no-rotation trials at washout as at training (480 trials), washout was often incomplete.

Incomplete washout can inadvertently magnify estimates of savings. Estimates of savings thus needs to take into account the extent of washout. We estimated percent adaptation for the test block as follows, where reach biases in the washout phase were estimated as the mean of the final 3 cycles of the washout block, similar to Haith et al. (2015).

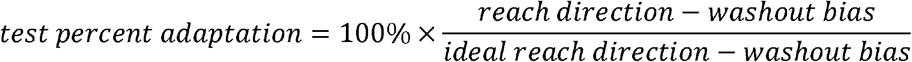

To evaluate savings (improved adaptation compared to naïve), we selected first 5 cycles (i.e., the first 40 trials) of the naïve and test block for comparisons using Welch’s t-tests (Delacre et al., 2017)(or Mann-Whitney U-tests if assumptions of normality were violated), as savings tends to be most evident at initial exposure/re-exposure to the rotation in visuomotor rotation paradigms. We note however that this window may miss important effects outside the first 5 cycles, particularly since adaptation is slower when task errors are removed or enforced (Leow et al., 2018; Kim et al., 2019). To avoid missing effects outside the first 5 cycles of the block, we additionally estimated performance in the entire adaptation block by splitting each 60-cycle adaptation block into an early phase (estimated as mean percent adaptation from the first 30 cycles) and a late phase (estimated as mean percent adaptation from the last 30 cycles). To compare naïve to test adaptation performance, we used Training (naïve, test) × Phase (early, late) ANOVAs on early and late phase percent adaptation, with greenhouse-geisser corrections applied where appropriate. For Experiment 1, test block adaptation was compared to naïve block adaptation within the same group of participants. For Experiment 2, test block adaptation with a certain task error manipulation was compared to naïve adaptation from a different group with equivalent task error manipulations. Specifically, the TrainStandardTE group and the TrainEnforcedTE group experienced no task errors at test, and thus were compared to the naïve no task error block from the TrainNoTaskErrors group. The TrainNoTaskErrors group experienced standard task errors at test and thus were compared to the naïve standard task error block in the TrainStandardTaskErrors group. Similarly, for Experiment 3, test block adaptation with standard task errors was compared with the naïve standard task error block in the TrainStandardTaskError group from Experiment 2. For Experiment 4, test block adaptation with short preparation time and standard task errors were compared with data from a separate control group who experienced the same short preparation time and standard task error conditions, but who were naïve to any training to reduce task errors or sensory prediction errors.

A common alternative measure of savings is to assess rate constants obtained by fitting the data to exponential functions. Rate constant analyses were not used here for the following reasons. For datasets where savings is evident as immediate adaptation in the first cycle upon perturbation exposure (Landi et al., 2011; Huberdeau et al., 2015), rate constants are typically small because the rapid initial adaptation is not captured by the fit. This can give the erroneous impression that savings is absent, and this situation was apparent in some of our data. When we tried to avoid this problem by fixing the fit parameter that reflects the y value when × = 0 as the mean reach direction in the immediately previous no-rotation cycle, the fits poorly characterized the data. These results agree with previous work demonstrating that exponential functions poorly represent individual learning curves, which often show abrupt step-like increases in performance (Gallistel et al., 2004).

Statistical analyses were performed with SPSS and JASP. Graphs were plotted with GraphPad Prism version 7.00 for Windows, GraphPad Software, La Jolla California USA, www.graphpad.com.

## Results

### Experiment 1: Task errors are important for savings

In Experiment 1, we asked whether savings would be present if adaptation was learnt in the absence of task errors, or with task errors that were enforced regardless of the participants’ actions. During training, all participants were exposed to a 30° rotation of cursor feedback. In the StandardTaskError condition, since the target was not moved within a trial, task errors were allowed to vary contingent upon participants’ responses to the cursor perturbation (Figure 1B). Task errors were enforced in the EnforcedTaskError condition by moving the target mid-movement so that the cursor always missed the target by 20-30° (Figure 1E). Task errors were removed in the NoTaskError condition by moving the target to align with the cursor trajectory mid-movement (Figure 1H). After initial exposure to the visuomotor rotation, behaviour was returned to the unadapted state by removing the cursor rotation in a washout phase. During this washout, we also employed the NoTaskError manipulation across all groups to prevent experience of task errors upon abrupt removal of the cursor rotation. At the test block (i.e., the second exposure to the visuomotor rotation) we applied the same task error manipulations that each group experienced when they were initially exposed to the perturbation.

Reach directions across all cycles for groups are shown in Figure 1B. Both removing task errors and enforcing task errors slowed adaptation compared to experiencing standard task errors that were contingent upon the corrective responses of the participant. One participant from the no task error group did not move towards the presented target in the cycles 41 to 47, which resulted in the variability that is apparent in the group average plot. The analysis outcomes were similar with and without this dataset.

Figure 2 shows percent adaptation in the second exposure to visuomotor rotation in comparison to naïve. With standard task errors, percent adaptation at test tended to improve compared to naive immediately after rotation onset (Figure 2A), similar to Huberdeau et al. (2015). This improvement was primarily evident in the first 8 trials of exposure to the cursor rotation (i.e., the first cycle), as adaptation in the first cycle improved at test compared to naive (naïve: 8.5+/−6.6%, test: 32.0+/−8.6%, cohen’s d=0.74). Improvements compared to naïve were marginal when estimated over the first 5 cycles (Figure 2B, naïve: 37.5+/− 3.9%; test: 53.4+/−8.6%, p= 0.055, cohen’s d= 0.618). Block × Phase (early adaptation, late adaptation) ANOVA comparing the naïve and the test block showed a non-significant main effect of block and no significant interactions with block.

**Figure 2.**
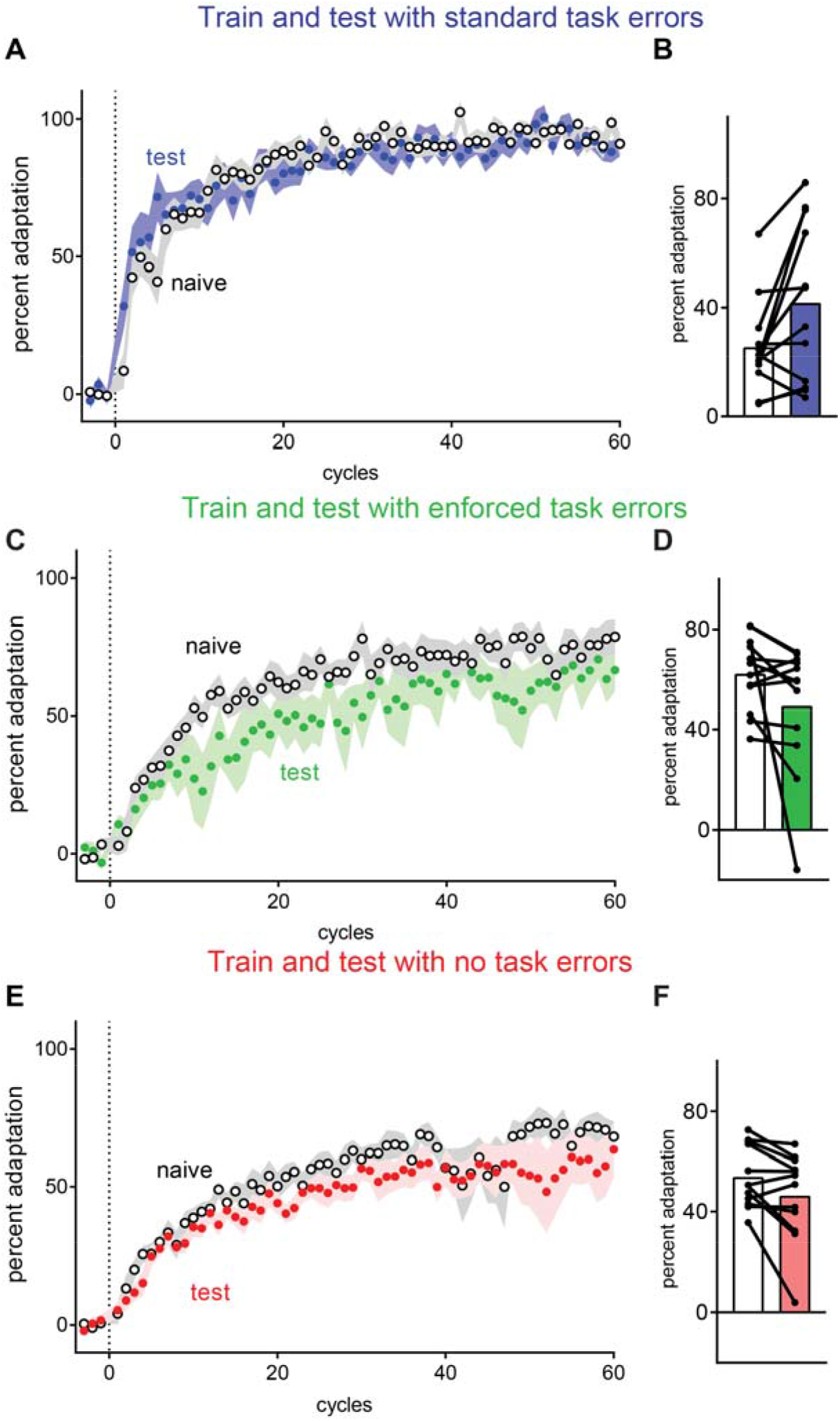
Experiment 1: Despite re-exposure to the same rotation, savings was absent with no task errors or enforced task errors. Cycle averaged percent adaptation (filled circles) compared to naïve (white circles) (A, C, E, error bars = SEM), and mean percent adaptation averaged across the first 5 cycles (B, D, F, error bars=95% CI).

With enforced task errors, percent adaptation in the first 5 cycles (Figure 2D) did not differ reliably between test (15.9+/−2.7%) and naïve (18.5+/−3.0%), t(11)=0.663, p=.521, cohen’s d=0.192. Percent adaptation at test showed a tendency to be almost worse compared to naive (Figure 2C), although Block × Phase (early adaptation, late adaptation) ANOVA comparing the naïve and the test block showed a non-significant main effect of block, F(1,11)=2.855, p=0.119, partial η-squared=0.2, and no reliable interactions with block.

With no task errors, percent adaptation in the first 5 cycles did not differ reliably from naïve (17.7+/−3.1%) to test (13.1+/−2.8%), t(11)=1.08, p=.303, cohen’s d =0.3. Analyses on the whole adaptation block showed that adaptation was worse at test than naïve, as Block × Phase ANOVA on the entire adaptation block showed a significant main effect of block, F(1,11)=5.95, p=0.033, partial η-squared=0.35, (mean of the entire naïve block, 53.0+/−3.6%, mean of the entire test block: 46.3+/− 5.1%).

Thus, despite exposure to the same cursor perturbation, and therefore previous experience of similar sensory prediction errors, savings was not evident in the groups that did not experience correctable task errors as a result of the cursor perturbation.

### Experiment 2: Task errors are required at encoding but not at retrieval

The absence of savings when perturbation-induced task errors were removed suggests some role of perturbation-induced task errors in savings. A few interpretations are possible. First, task errors might act as a retrieval cue to trigger the memory that is responsible for savings (Huberdeau et al., 2015). Second, task errors might be necessary to encode a memory that is responsible for savings. Third, task errors might be necessary both at encoding and at retrieval for savings: savings can only occur when previously experienced task errors are revisited. We dissociated these possibilities in Experiment 2. Task errors were manipulated either at training or at test to identify whether savings requires prior experience of task errors during first exposure to a perturbation (when a memory is first “encoded”) or when the perturbation is re-encountered (when a memory is “retrieved”): see Figure 1B. A TrainStandardTaskError group (n=12, 6CW, 6CCW) was deprived of task errors at test (target was shifted mid-movement so that the cursor always hit the target), but were provided standard task errors at training (i.e. no target shifts): absence of savings here would suggest that task errors are necessary as a retrieval cue for savings. A TrainNoTaskError group (n=12, 6CW, 6CCW) was deprived of task errors at training, but experienced standard task errors at test (target did not move mid-movement): absence of savings here would suggest that the task errors are not required as a retrieval cue, but are a necessary component to encoding a memory that results in savings. Does savings result from the experience of task errors alone, or does savings require learning to correct for task errors? To test this, a TrainEnforcedTaskError group (n=12,6CW, 6CCW) were provided with enforced task errors at training (target always moves away from the cursor mid-trial, such that they could never succeed in reducing task errors), and were tested for savings in the absence of task errors. After training, all groups encountered 6 no-feedback cycles and 60 no-rotation washout cycles with no task error, and then re-encountered the same cursor rotation as they experienced at training.

Figure 1D shows that the effects of the task error manipulation at training in Experiment 2 were consistent with Experiment 1. To evaluate savings, test performance was compared to the naive adaptation of another group who experienced the same task error conditions. Specifically, the no task error test phase from the TrainStandardTaskError and the TrainEnforcedTaskError groups was compared to the no task error training phase of the TrainNoTaskError group (Figure 3A & C). The standard task error test phase from the TrainNoTaskError group was compared to the standard task error training phase of the TrainStandardTaskError group (Figure 3E).

**Figure 3.**
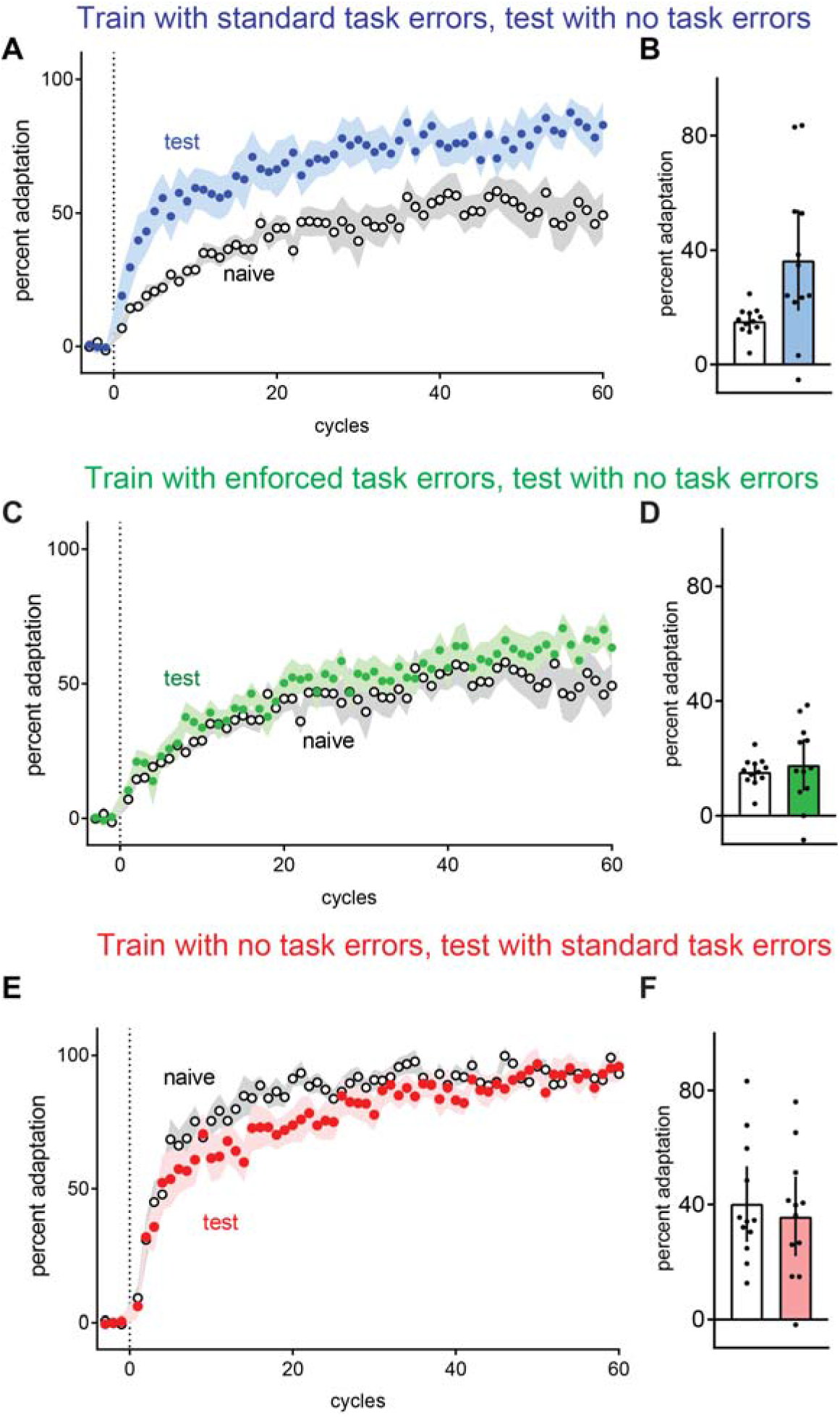
Experiment 2 showed that savings requires a history of adaptation to task errors. During initial exposure to the cursor rotation (training), participants experienced task errors (either standard task errors, enforced task errors, or no task error). At re-exposure to the cursor rotation (test), participants who experienced task errors at training were now deprived of task errors, whereas participants who were deprived of task errors at training were now provided task errors. Clear symbols and white bars indicate naïve adaptation. **(B)**. Even without task errors at test, a history of standard task errors at training improved subsequent adaptation, (greater percent adaptation in A & B). In contrast, adaptation was unimproved in the group who experienced a history of enforced task errors that could not be corrected (C&D). Adaptation was also unimproved without a history of task errors at training, despite the presence of task errors at test (E & F). For B, D, F, error bars=95% CI. All other error bars are SEM.

**Figure 4.**
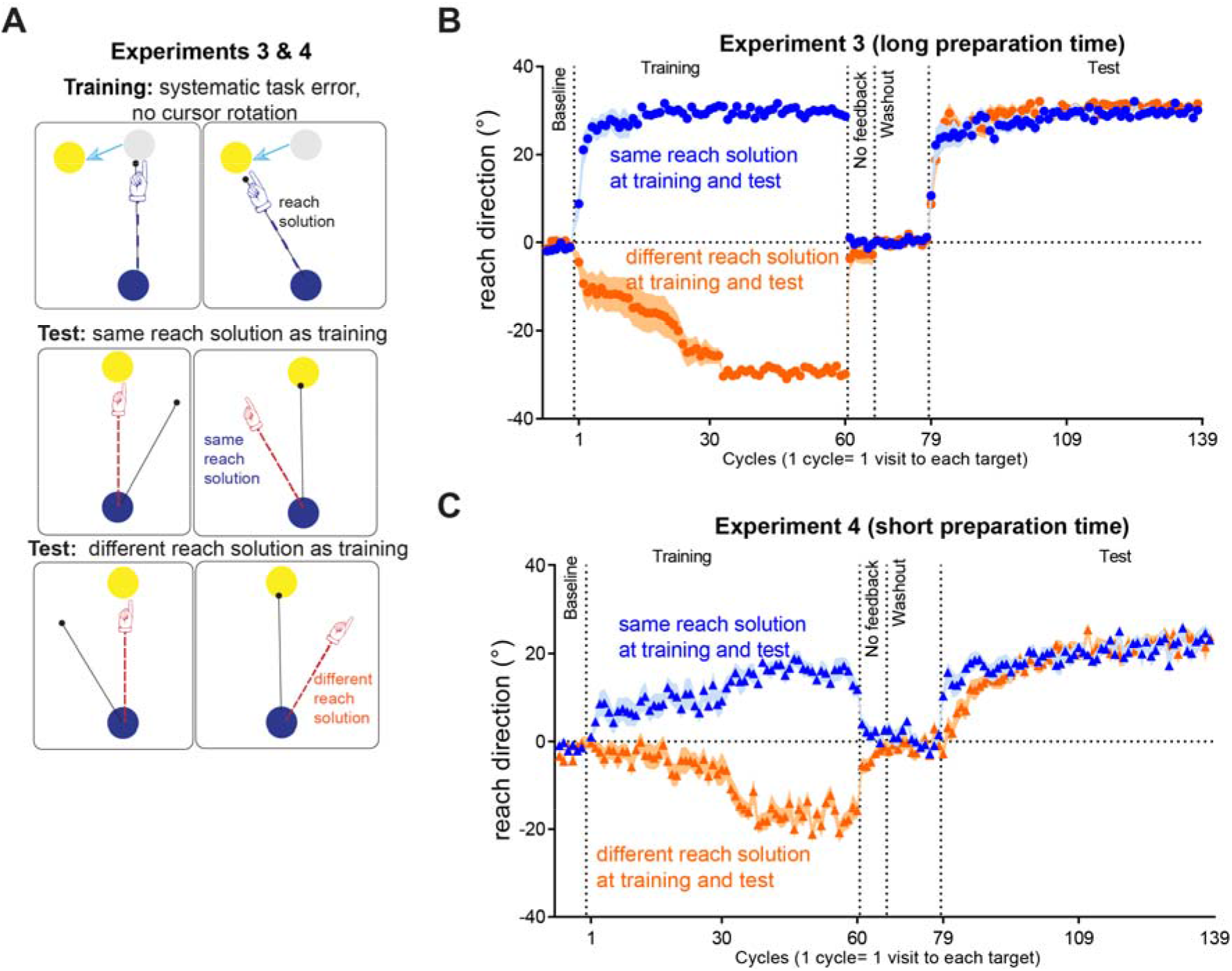
Experiments 3 and 4 test if learning to correct for task errors alone can evoke savings when subsequently exposed to a (novel) visuomotor rotation. At training, participants learnt to counteract task errors resulting from the target moving away from its original position by 30° at mid-movement (4cm into the 9cm start-target distance). To hit the target, participants had to to move 30° away from the presented target. After training, participants rapidly disengaged this learning upon instructions that all disturbances had been removed. At test, participants encountered a 30° cursor rotation for the first time. The Same group had the same reach solution at test as at training (i.e., the cursor rotation could be counteracted by the target movement direction during training). The Different group had a directionally opposite reach solution at training and at test, (the cursor rotation was in the same direction as the target movement direction during training). Experiment 4 replicates the design of Experiment 3, except that we limited the amount of time available to prepare movement.

Figure 3A shows that in the TrainStandardTaskError group, standard task errors at training resulted in better-than-naïve adaptation at test, even in the absence of task errors at test, as evidenced in better adaptation at the first 5 cycles at test (36.5+/−8.0%) compared to naive (15.2+/−1.5%), t(11.7)= 2.61, p= 0.016, cohen’s d=1.065. Similarly, Group (TrainStandardTaskError, Naïve) × Phase (Early Adaptation, Late Adaptation) ANOVA showed a significant main effect of Group: F(1,22)=10.211, p= .004, partial eta-squared =.317. Thus, a history of task errors improved re-adaptation to a cursor rotation even when the perturbation did not cause any task errors upon second exposure.

In contrast, the TrainEnforcedTaskError group failed to improve adaptation compared to naïve (Figure 3C). Adaptation in the first 5 cycles did not differ reliably at test (17.6+/−4.1%) compared to naïve (15.2+/−1.4%), t(13.7) = 0.561, p = 0.584, cohen’s d = 0.229. Group (TrainEnforcedTaskError, Naïve) × Phase ANOVA on the whole adaptation block similarly showed no reliable main effect of Group, F(1,22) = 1.20, p=.285, partial eta-squared=.052, and a non-significant Phase × Group interaction, F(1,22) = 0.69, p = 0.413, partial η-squared = 0.03. Enforcing task errors at training thus did not appear to improve adaptation compared to naïve when tested without task errors. Thus, merely experiencing task errors, without learning to correct for those task errors, was not sufficient to evoke subsequent savings.

In the TrainNoTaskError group (Figure 3F), adaptation in the first 5 cycles did not differ reliably at test (35.9+/−6.3%) compared to naïve (40.3+/−6%), t(21.9) = 0.499, p = 0.623, cohen’s d = 0.204. This lack of improvement compared to naïve was shown throughout the entire adaptation block (Figure 2I), as Group (TrainNoTaskError, Naïve) × Phase (Early, Late Adaptation) ANOVA showed a non-reliable main effect of group, F(1,22) = 1.38, p = 0.25, partial η-squared = 0.05, and a non-reliable Phase × Group interaction, F(1,22) = 3.47, p = 0.08, partial η-squared = 0.13. Thus, depriving participants of task errors when they were first exposed to the cursor rotation at training resulted in no savings despite the subsequent presence of standard task errors at test.

Thus, a history of adapting movements to correct task errors appears necessary to encode learning that improves adaptation to a previously experienced visuomotor rotation. The presence of task errors appears unnecessary to retrieve this learning.

### Experiment 3: Learning to correct for task errors is sufficient to evoke savings

Recent work suggests that savings in visuomotor rotation primarily reflects the deliberate application of a strategy, where participants explicitly re-aim to the one side of a target to counteract the rotation of cursor feedback (Haith et al., 2015; Morehead et al., 2015). This view considers the role of implicit adaptation to sensory prediction errors as secondary to the role of strategy in savings, and would interpret the presence/absence of savings with task errors in Experiments 1 and 2 to be because task errors provoke the formation of explicit strategies. An alternative view is that task errors alter the sensitivity to sensory prediction errors (Leow et al., 2018; Kim et al., 2019), and increased sensitivity to these errors produce savings (Herzfeld et al., 2014). We cannot dissociate between the two alternative explanations based on data from Experiments 1 and 2 alone because task errors here were not wholly independent of sensory prediction errors; the task error manipulations were always made in the presence of a perturbation of the hand-cursor relationship (which therefore induced sensory prediction errors). Thus, we next examined how learning to correct for task errors alone, in the absence of sensory prediction errors (i.e., in the absence of any perturbation of the hand-cursor relationship), affected subsequent adaptation to a visuomotor rotation.

In Experiment 3, we did not perturb the cursor at training, but enforced systematic task errors that could be counteracted by a re-aiming strategy: the target always moved away by 30° from the original target location mid-movement: participants could correct these task errors by re-aiming 30° away from the original target (see Figure 3a). For one group of participants, the reach solution needed to hit a given target after it jumped mid-movement was the same reach solution needed to counteract the cursor rotation in the test block (**Same**, n=12, 6 CW, 6 CCW). For example, if the target jump at training was 30° counterclockwise, then the cursor rotation at test was 30° clockwise (thus requiring a counterclockwise compensatory hand movement). To test whether this learning is flexible enough to accommodate a different reach solution, we had another group of naïve participants (**Different**, n=12, 6 CCW, 6CW), where the reach direction required to hit targets at training was opposite to that at test. Pilot testing revealed substantial individual differences in how quickly participants developed a strategy to re-aim at training. Thus, if the experimenter observed that participants had yet to show successful re-aiming by trial 180 of the 480-trial training block, the experimenter explicitly instructed participants that a strategy may be needed to hit the target. This explicit instruction was required in 2 of the 12 participants in the Same group, and 6 of the 12 participants in the Different group. Here, the instruction was given without exposure to the cursor perturbation but after exposure to the *task* error. At the test block, no instructions about re-aiming were provided. To quantify savings, we compared percent adaptation at test to naïve controls who experienced similar task error manipulations (i.e., the naïve adaptation block from the group who experienced standard task errors at training in Experiment 2).

Figure 4B shows reach directions in all cycles. After instructing participants that the task error manipulation had been removed, reach directions reverted rapidly back to baseline in the no-feedback block. This illustrates that the re-aiming response can be switched off immediately upon instruction, similar to previous work (e.g., Mazzoni and Krakauer, 2006; Benson et al., 2011; Taylor and Ivry, 2011; Schween et al., 2014; Savoie et al., 2018),

Despite being naïve to the cursor rotation, improved adaptation was evident when the reach solution at test was the same as training (Figure 5A &B), as better adaptation at test was evident in the first 5 cycles at test (69.1+/−8.7%) compared to naïve (40.3+/−6%), t(19.5) = 2.722, p = 0.013, cohen’s d = 1.111. This effect was primarily limited to the first 5 cycles, as analyses on the entire adaptation block via Group × Phase (Early, Late Adaptation) ANOVA showed a main effect of Group, F(1,22) = 2.82, p = 0.107, partial η-squared = 0.11, and an Phase × Group interaction, F(1,22) = 1.1, p = 0.304, partial η-squared = 0.04. Similarly, improved adaptation was evident when the reach solution at test was opposite to that at training (Figure 3E), as evident in better adaptation in the first 5 cycles at test (72.2+/−6.7%) compared to naïve (40.3+/−6%), t(21.7) = 3.558, p=0.002, cohen’s d=1.453. Group × Phase (Early, Late Adaptation) ANOVA showed a main effect of Group, F(1,22) = 8.11, p = 0.00934, partial η-squared = 0.26. The Phase × Group interaction was not reliable, F(1,22) = 1.75, p = 0.199, partial η-squared = 0.07. We note that this result stands in contrast to a previous finding of interference from previously learning an incongruent reach solution (Zach et al., 2005), but in that study, no washout nor task instruction nor change in task context was given to participants to disengage the previous reach solution: interference there might be because participants unwittingly continued behaving as before.

**Figure 5.**
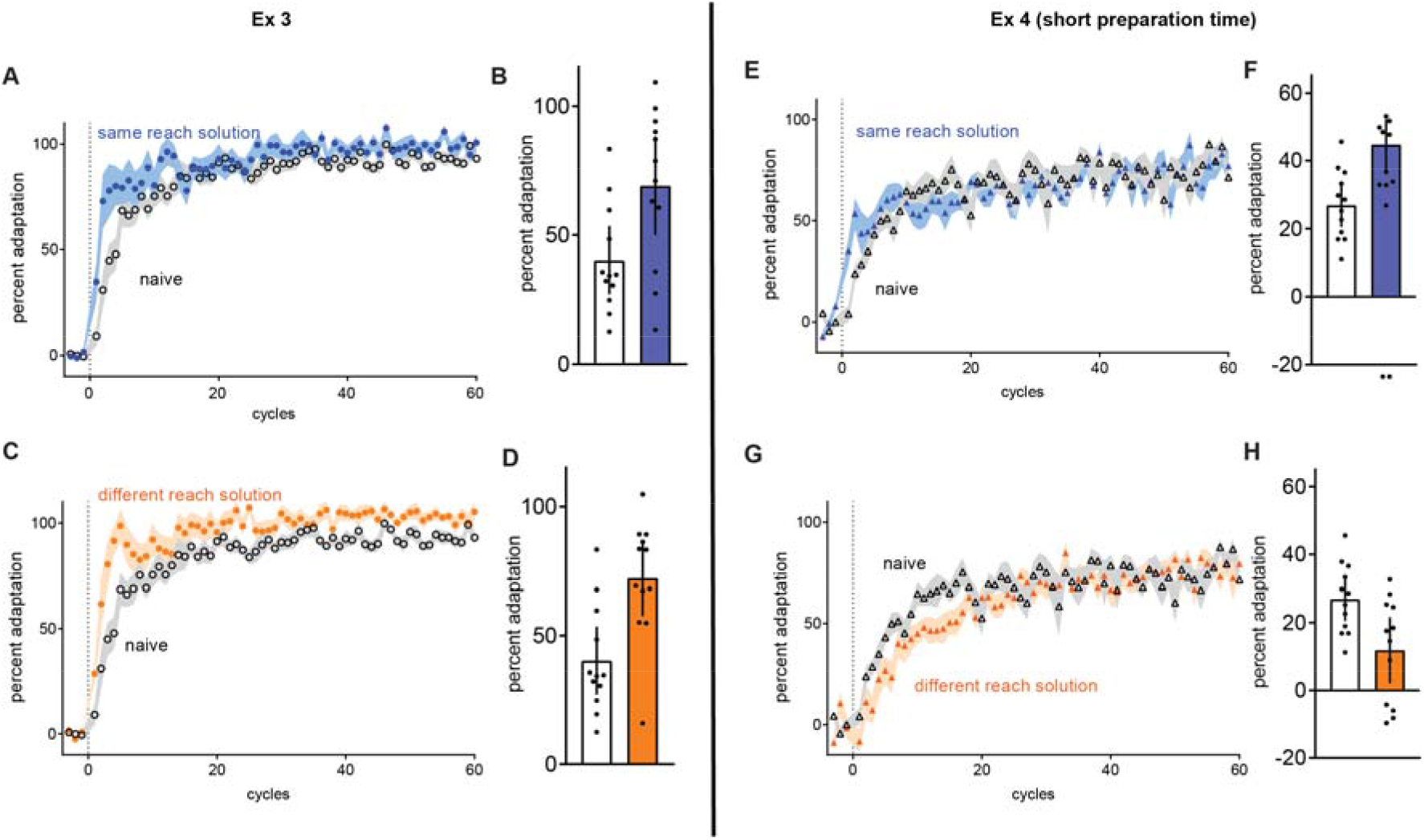
Adaptation to visuomotor rotation after learning to correct for task errors with veridical cursor feedback (filled symbols), compared to naïve (clear symbols) in Experiments 3 and 4. Cycle-averaged percent adaptation during test (filled symbols) compared to naïve (clear symbols) for the same reach solution (A) and the different reach solution (C). Bar graphs show mean percent adaptation averaged over the first 5 cycles (B & D). Savings was evident in more adapted reaches compared to naïve (clear symbols) for both the Same group (A & B) and the Different group (C & D). Experiment 4 replicates the design of Experiment 3, except that participants had limited preparation time for each movement. Savings was evident in more adapted reaches compared to naïve (clear symbols) for the ShortSame group (E & F). The ShortDifferent group (G & H) showed worse adaptation compared to naïve.

Hence, previous learning to counteract task errors was sufficient to improve subsequent adaptation to a visuomotor rotation, even when participants were naïve to perturbation-induced sensory prediction errors. This learning was flexible: it elicited savings even when the reach solution required to hit a given target at training was different from the reach solution required to hit that target at test.

### Experiment 4: Suppressing goal-directed behaviour by reducing movement preparation time

We next explored the mechanisms by which learning to compensate for task errors can improve subsequent visuomotor rotation adaptation. Task error correction likely relies on strategic, goal-directed processes that demand cognitive resources, because error compensation is reduced by manipulations that limit cognitive resources, such as time-constraints or dual-tasking (Taylor and Thoroughman, 2007, 2008; Galea et al., 2010; Malone and Bastian; Fernandez-Ruiz et al., 2011; Anguera et al., 2012; Haith et al., 2015; Leow et al., 2017). In visuomotor rotation paradigms, these observations are consistent with the notion that mental rotation of a movement plan at a specified angle away from a target is computationally expensive, and requires time (Georgopoulos and Massey, 1987; Pellizzer and Georgopoulos, 1993; Bhat and Sanes, 1998). Does savings solely result from goal-directed cognitive strategies (Morehead et al., 2015)? To test this, we examine if savings occurred when these time-consuming cognitive processes were suppressed by limiting preparation time. We replicated the design of Experiment 3, except that participants were required to move within a short preparation time of 250ms after the time of target presentation throughout all trials. We used the timed-response paradigm described in the methods for all the previous experiments, but instead of presenting the target at 1000ms before the imperative to move, in experiment 4 we presented the target 250ms before the imperative to move (Leow et al., 2017). There were two conditions: **ShortSame** (n=12, 6 CW, 6 CCW), where the reach solution is the same at training and at test, and **ShortDifferent**, (n=12, 6 CW, 6 CCW), where the reach solution is opposite at training and at test. Explicit instructions to re-aim were necessary in 4 of the 12 participants in the ShortSame group, and 8 of the 12 participants in the Different group.

At training, despite the preparation time constraints, participants did learn to compensate for task errors (Figure 4B), as percent adaptation was larger at the late phase of the training block than the early phase, as shown by a significant main effect of Phase, F(1, 23) = 57.1, p=1.1e−7, partial eta-squared=0.7. However, the extent of compensation at training (estimated as the last 10 cycles) was less complete with short preparation time in Experiment 4 (ShortSame: 51.5+/−3.5%, ShortDifferent: 54.4+/−4.9%) than with long preparation time in Experiment 3 (Same 98.9+/−1.8%, Different: 97.4+/−2.5%), as ReachSolutionDirection (Same, Different) × Preparation Time (Short, Long) showed a significant main effect of preparation time F(1, 44) = 179.3, p = 4.0e-17, partial eta-squared = 0.8.

We compared test block adaptation to a naïve control group tested under similar preparation time and task error conditions (ShortNaive, n=12, 6 CW, 6 CCW). In the ShortSame group who trained with the same reach solution at test and training, adaptation at test was better than naïve (Figure 4C), as mean of the first 5 cycles of the test block, 44.7+/−3.4% was better than naïve, 26.9+/−2.9%, t(21.6)= 3.964, p=6.581e_-4, cohen’s d=1.618). Improved adaptation compared to naive was primarily concentrated in the first 5 cycles, as examining the entire adaptation block with a Group × Phase (Early, Late adaptation) ANOVA yielded an non-significant effect of Group F(1,22) = 0.01, p = 0.898, partial η-squared = 0, and a non-significant Group × Phase interaction, F(1,22) = 0.17, p = 0.679, partial η-squared = 0.

When the reach solution at test differed from that at training in the ShortDifferent group, test performance was ***worse*** than naïve (Figure 4E), as shown in worse percent adaptation in the first 5 cycles at test (11.8+/−4.4%) compared to naïve (26.9+/−2.9%), t(19.2)= 2.839, p= 0.010, cohen’s d = 1.159. This was despite the fact that participants could obviously disengage the previously learnt reach solution with instruction (Figure 4B). Group × Phase (Early, Late adaptation) on the entire adaptation block showed a Group × Phase interaction, F(1,22) = 9.59, p = 0.00524, partial η-squared = 0.3, as worse adaptation tended to occur in the early phase (the first 30 cycles) (see Figure 4E).

## Discussion

Here, we demonstrate a fundamental role of adaptation to task errors in savings: a hallmark of latent sensorimotor memory. Savings was absent without prior correction of task errors (Experiments 1 & 2). Prior correction of task errors, even without prior exposure to sensory prediction errors, was also sufficient to elicit savings (Experiments 3 & 4). Thus, task errors that prompt adaptive responses can affect subsequent adaptation to never-before encountered sensorimotor perturbations. Experiments 3 and 4 further demonstrate that these sensorimotor memories can rely on distinct processes: (1) a time-consuming process that is flexible enough to facilitate corrective responses in the opposite direction; (2) a rapidly-accessible process that can be volitionally disengaged, but is not sophisticated enough to produce a novel corrective response.

### Savings require prior adaptation to task errors

Despite extensive study, exactly what mechanisms underpin savings remain unresolved. One influential theory suggests that a history of errors increases sensitivity to those errors, improving learning when re-encountering familiar errors (Herzfeld et al., 2014). Sensorimotor perturbations typically evoke both SPEs and task errors, but it was previously unclear how these errors contribute to sensorimotor memories that improve learning (Orban de Xivry and Lefevre, 2015; Leow et al., 2016). Here we show that, at least in visuomotor rotation, a history of adaptation to task errors is crucial to encode memories that improve subsequent adaptation. Uncorrectable task errors (i.e., enforced task errors that occurred regardless of behaviour) failed to evoke savings. Furthermore, task errors need not be present upon re-experiencing the perturbation, contradicting the idea that task errors act as a retrieval cue to trigger savings (Huberdeau et al., 2015). We note however that the absence of a measurable contribution of SPEs to savings in visuomotor rotation might not mean zero contribution of SPEs to all forms of persistent sensorimotor memories.

### Savings do not require a history of sensory prediction errors

When task errors were absent in Experiments 1 and 2, the lack of savings despite abundant prior exposure to SPEs suggests that the task errors and not SPEs are compulsory for savings. One possibility is that task errors drive deliberate corrective responses during adaptation, and that savings results from faster reselection of previously learnt actions (Haith et al., 2015; Huberdeau et al., 2015; Morehead et al., 2015). If true, then deliberate correction of task errors alone in the absence of SPEs should be sufficient for savings. We tested this in Experiment 3 by imposing a systematic task error at training (targets always jumped mid-movement by 30°) without perturbing hand position feedback. Learning to correct these task errors improved subsequent naïve adaptation to the 30° rotation of hand position feedback. Thus, learning to counteract task errors in the absence of a sensorimotor perturbation was enough to improve adaptation to a never-before encountered perturbation. However, savings was not merely due to faster reselection of previously learned actions, since we also observed adaptation benefits for different actions.

### How might correction of task errors lead to savings?

What is encoded when correcting task errors? In Experiment 3, we found that adaptive responses to task errors can be volitionally disengaged, but this learning was retained in latent form to affect subsequent adaptation. Thus, this form of learning seems fundamentally distinct from adaptive responses to SPEs, which are expressed obligatorily after perturbation removal (Izawa and Shadmehr, 2011), indicating a remapping between motor intent and its predicted sensory outcomes. In contrast, the contextually flexible response to task errors might arise from an earlier component of the sensory-to-motor transformation: a mapping between the behavioural goal and the motor plan selected to achieve it (Crawford et al., 2011).

Given that we can instruct participants to deliberately use re-aiming strategies (e.g., Mazzoni and Krakauer, 2006), it is perhaps unsurprising that task errors led to savings by prompting acquisition of a re-aiming strategy. Indeed, one view is that savings results solely from deliberate strategy (Morehead et al., 2015). This view is supported by the absence of savings (Haith et al., 2015) when time pressure is used to suppress time-consuming mental rotation processes required to re-aim accurately away from the presented target (e.g., Georgopoulos and Massey, 1987). Although recent findings show savings despite time pressure with repeated exposure to opposing visuomotor rotations (Huberdeau et al., 2017), savings in that study might have resulted partly from improvements in the residual capacity to re-aim away from a target at short latencies, when the range of required responses is small and predictable (Leow et al., 2017; McDougle and Taylor, 2019).

But is the capacity to employ mental rotation the sole contributor to savings in visuomotor rotation? We explored this in Experiment 4 by replicating Experiment 3, but limiting preparation time. Targets were randomly presented in a wide spatial array, reducing predictability of the required response. With time-pressure, reaches only partially compensated for the rotation at training (only ≈50% percent adaptation despite 480 training trials), consistent with previous work (Leow et al., 2017; McDougle and Taylor, 2019). At test, adaptation was better-than-naive when the reach solution at test matched that at training, and *worse*-than-naïve when the reach solutions were opposite at training and test. Thus, although time-pressure increases the difficulty of mental rotation, it might not prevent the formation of associations between a given stimulus and the required response (i.e., stimulus-response associations). After washout, these latent stimulus-response associations could have been re-expressed, eliciting savings when reach solutions at training matched that at test, and interference when the reach solutions differed. Intriguingly, participants re-engaged a now-maladaptive reach solution even when they were obviously able to immediately disengage it at washout under similar time-pressure. Perhaps the learnt reach solution was triggered by some contextual cue (e.g., experiencing errors to the side of targets), and this pre-potent response to the trigger was poorly inhibited under time pressure. This is consistent with the tendency to re-express learnt stimulus-response associations even when they result in undesired outcomes (Dolan and Dayan, 2013). Alternatively, interference might result from prior training in mental rotation in the opposite direction (Sack et al., 2007). One clue to identify the more likely explanation comes from findings of deficient savings (Marinelli et al., 2009; Bedard and Sanes, 2011; Leow et al., 2012; Leow et al., 2013) and anterograde interference in Parkinson’s disease (Leow et al., 2013). Parkinson’s disease patients are better than controls when adapting to a rotation opposite to that previously experienced (Leow et al., 2013). This seems unlikely to result from superior capacity to disengage effects of previous mental rotation in an opposite direction. Deficient learning of stimulus-response associations in Parkinson’s disease (Knowlton et al., 1996; Foerde, 2018) seems to be the more likely culprit. Thus, we interpret our data as evidence for the contribution of stimulus-response associations to sensorimotor memories.

The insight that stimulus-response associations contribute to sensorimotor memories has implications for work examining how the brain stores and retrieves sensorimotor memories. Associative learning relies on corticostriatal circuits (Toni et al., 2001; Yin and Knowlton, 2006; Rueda-Orozco and Robbe, 2015). During adaptation, learning stimulus-response associations might similarly rely on corticostriatal circuits, consistent with the abovementioned work in Parkinson’s disease, and with imaging work implicating the basal ganglia in adaptation (Shadmehr and Holcomb, 1997; Shadmehr and Holcomb, 1999; Diedrichsen et al., 2005; Seidler et al., 2006). In particular, target-errors activate the striatum (Diedrichsen et al., 2005) and area 7 of the parietal cortex (Diedrichsen et al., 2005; Inoue and Kitazawa, 2018). We speculate that task errors trigger an assessment of task state by the basal ganglia, prompting remapping of the relationship between goals and movement plans by parietal cortex area 7. Desirable task outcomes (e.g., reduced task error) might trigger the formation of stimulus-response associations, which is known to concurrently alter activity in the dorsal premotor cortex and parts of the putamen receiving inputs from the dorsal premotor cortex (Brasted and Wise, 2004). Thus, the characteristic changes in motor/dorsal premotor cortex preparatory activity seen in sensorimotor adaptation (e.g., Li et al., 2001; Paz et al., 2003; Mandelblat-Cerf et al., 2011; Perich et al., 2018; Zhou et al., 2019) might partly reflect how movement plans are affected by stimulus-response associations, as well as by implicit remapping in response to sensory prediction errors that occur via changes in functional connectivity between the cerebellum, area 5 of the parietal cortex, and motor cortex (Tanaka et al., 2009; Haar et al., 2015; Inoue and Kitazawa, 2018). Notably, our current data suggest that persistent sensorimotor memories that improve relearning rely primarily on cortico-striatal circuits, rather than cortico-cerebellar circuits. Future work involving neural recordings that dissociate task and sensory prediction errors is required to test this prediction.

## Summary

Our results show that failures to attain movement goals, or task errors, are a fundamental driver of latent memories in sensorimotor adaptation. Sensorimotor memories are influenced not only by time-consuming, goal-directed processes (such as mental rotation), but also by rapidly-accessible, stimulus-response associations. The data demonstrate the richness of behavioural responses to sensorimotor perturbations, and suggest that future efforts to characterise the neural mechanisms underpinning persistent sensorimotor memories should benefit from explicit dissociation of the distinct behavioural processes involved.

## Acknowledgments

The study was funded by an Australian Research Council Discovery Project Grant DP180103081. We would like to thank Aya Uchida, Tess Stevenson, and Tamara Spingler for assistance with data collection.

## References

Anguera JA, Bernard JA, Jaeggi SM, Buschkuehl M, Benson BL, Jennett S, Humfleet J, Reuter-Lorenz PA, Jonides J, Seidler RD (2012) The effects of working memory resource depletion and training on sensorimotor adaptation. Behav Brain Res 228:107–115.

Avraham G, Keizman M, Shmuelof L (2019) Environmental Consistency Modulation of Error Sensitivity During Motor Adaptation is Explicitly Controlled. bioRxiv:528752.

Bedard P, Sanes JN (2011) Basal ganglia-dependent processes in recalling learned visual-motor adaptations. Exp Brain Res 209:385–393.

Benson BL, Anguera JA, Seidler RD (2011) A spatial explicit strategy reduces error but interferes with sensorimotor adaptation. J Neurophysiol 105:2843–2851.

Bhat RB, Sanes JN (1998) Cognitive channels computing action distance and direction. J Neurosci 18:7566–7580.

Bostan AC, Strick PL (2018) The basal ganglia and the cerebellum: nodes in an integrated network. Nat Rev Neurosci 19:338–350.

Brasted PJ, Wise SP (2004) Comparison of learning related neuronal activity in the dorsal premotor cortex and striatum. European Journal of Neuroscience 19:721–740.

Braun DA, Aertsen A, Wolpert DM, Mehring C (2009) Motor task variation induces structural learning. Curr Biol 19:352–357.

Crawford JD, Henriques DYP, Medendorp WP (2011) Three-dimensional transformations for goal-directed action. Annual review of neuroscience 34:309–331.

Cunningham HA (1989) Aiming error under transformed spatial mappings suggests a structure for visual-motor maps. J Exp Psychol Hum Percept Perform 15:493–506.

de Brouwer AJ, Albaghdadi M, Flanagan R, Gallivan JP (2017) Gaze Behaviour During Sensorimotor Adaptation Parcellates the Explicit and Implicit Contributions to Learning. J Neurophysiol.

Delacre M, Lakens D, Leys C (2017) Why Psychologists Should by Default Use Welch’s t-test Instead of Student’s t-test. International Review of Social Psychology 30.

Diedrichsen J, Hashambhoy Y, Rane T, Shadmehr R (2005) Neural correlates of reach errors. J Neurosci 25:9919–9931.

Dietz V, Zijlstra W, Duysens J (1994) Human neuronal interlimb coordination during split-belt locomotion. Exp Brain Res 101:513–520.

Dolan RJ, Dayan P (2013) Goals and habits in the brain. Neuron 80:312–325.

Ebbinghaus H (1964) Memory: A contribution to experimental psychology. Memory: A Contribution to Experimental Psychology.

Favilla M, De Cecco E (1996) Parallel direction and extent specification of planar reaching arm movements in humans. Neuropsychologia 34:609–613.

Fernandez-Ruiz J, Wong W, Armstrong IT, Flanagan JR (2011) Relation between reaction time and reach errors during visuomotor adaptation. Behav Brain Res 219:8–14.

Foerde K (2018) What are habits and do they depend on the striatum? A view from the study of neuropsychological populations. Current opinion in behavioral sciences 20:17–24.

Galea JM, Sami SA, Albert NB, Miall RC (2010) Secondary tasks impair adaptation to step- and gradual-visual displacements. Exp Brain Res 202:473–484.

Gallistel CR, Fairhurst S, Balsam P (2004) The learning curve: implications of a quantitative analysis. Proc Natl Acad Sci U S A 101:13124–13131.

Georgopoulos AP, Massey JT (1987) Cognitive Spatial-Motor Processes .1. The Making of Movements at Various Angles from a Stimulus Direction. Experimental Brain Research 65:361–370.

Gonzalez Castro LN, Hadjiosif AM, Hemphill MA, Smith MA (2014) Environmental consistency determines the rate of motor adaptation. Curr Biol 24:1050–1061.

Haar S, Donchin O, Dinstein I (2015) Dissociating visual and motor directional selectivity using visuomotor adaptation. Journal of Neuroscience 35:6813–6821.

Hadjiosif A, Smith M (2013) Savings is restricted to the temporally labile component of motor adaptation. In: Translational and Computational Motor Control. Washington DC.

Haith AM, Huberdeau DM, Krakauer JW (2015) The influence of movement preparation time on the expression of visuomotor learning and savings. J Neurosci 35:5109–5117.

Heffley W, Song EY, Xu Z, Taylor BN, Hughes MA, McKinney A, Joshua M, Hull C (2018) Coordinated cerebellar climbing fiber activity signals learned sensorimotor predictions. Nat Neurosci 21:1431–1441.

Herzfeld DJ, Vaswani PA, Marko MK, Shadmehr R (2014) A memory of errors in sensorimotor learning. Science 345:1349–1353.

Howard IS, Ingram JN, Wolpert DM (2009) A modular planar robotic manipulandum with end-point torque control. J Neurosci Methods 181:199–211.

Huang VS, Haith A, Mazzoni P, Krakauer JW (2011) Rethinking motor learning and savings in adaptation paradigms: model-free memory for successful actions combines with internal models. Neuron 70:787–801.

Huberdeau DM, Haith AM, Krakauer JW (2015) Formation of a long-term memory for visuomotor adaptation following only a few trials of practice. J Neurophysiol 114:969–977.

Huberdeau DM, Krakauer JW, Haith AM (2017) Practice induces a qualitative change in the memory representation for visuomotor learning. bioRxiv:226415.

Inoue M, Kitazawa S (2018) Motor Error in Parietal Area 5 and Target Error in Area 7 Drive Distinctive Adaptation in Reaching. Curr Biol 28:2250–2262 e2253.

Izawa J, Shadmehr R (2011) Learning from sensory and reward prediction errors during motor adaptation. PLoS Comput Biol 7:e1002012.

Kim HE, Parvin DE, Ivry RB (2019) The influence of task outcome on implicit motor learning. Elife 8:363606.

Knowlton BJ, Mangels JA, Squire LR (1996) A neostriatal habit learning system in humans. Science 273:1399–1402.

Kostadinov D, Beau M, Pozo MB, Hausser M (2019) Predictive and reactive reward signals conveyed by climbing fiber inputs to cerebellar Purkinje cells. Nat Neurosci 22:950–962.

Krakauer JW, Carmichael ST (2017) Broken Movement: The Neurobiology of Motor Recovery After Stroke: MIT Press.

Landi SM, Baguear F, Della-Maggiore V (2011) One week of motor adaptation induces structural changes in primary motor cortex that predict long-term memory one year later. J Neurosci 31:11808–11813.

Leow L-A, De Rugy A, Loftus AM, Hammond G (2013) Different mechanisms contributing to savings and anterograde interference are impaired in Parkinson’s. Frontiers in Neuroscience:108.

Leow L-A, De Rugy A, Marinovic W, Riek S, Carroll TJ (2016) Savings for visuomotor adaptation require prior history of error, not prior repetition of successful actions. Journal of Neurophysiology 116:1603–1614.

Leow LA, Loftus AM, Hammond GR (2012) Impaired savings despite intact initial learning of motor adaptation in Parkinson’s disease. Exp Brain Res 218:295–304.

Leow LA, Gunn R, Marinovic W, Carroll TJ (2017) Estimating the implicit component of visuomotor rotation learning by constraining movement preparation time. J Neurophysiol 118:666–676.

Leow LA, Marinovic W, de Rugy A, Carroll TJ (2018) Task errors contribute to implicit aftereffects in sensorimotor adaptation. Eur J Neurosci 48:3397–3409.

Li CS, Padoa-Schioppa C, Bizzi E (2001) Neuronal correlates of motor performance and motor learning in the primary motor cortex of monkeys adapting to an external force field. Neuron 30:593–607.

Maeda RS, McGee SE, Marigold DS (2018) Long-term retention and reconsolidation of a visuomotor memory. Neurobiol Learn Mem 155:313–321.

Malone LA, Bastian AJ (2010) Thinking about walking: effects of conscious correction versus distraction on locomotor adaptation. J Neurophysiol 103:1954–1962.

Mandelblat-Cerf Y, Novick I, Paz R, Link Y, Freeman S, Vaadia E (2011) The neuronal basis of long-term sensorimotor learning. J Neurosci 31:300–313.

Marinelli L, Crupi D, Di Rocco A, Bove M, Eidelberg D, Abbruzzese G, Ghilardi MF (2009) Learning and consolidation of visuo-motor adaptation in Parkinson’s disease. Parkinsonism Relat Disord 15:6–11.

Mazzoni P, Krakauer JW (2006) An implicit plan overrides an explicit strategy during visuomotor adaptation. J Neurosci 26:3642–3645.

McDougle SD, Taylor JA (2019) Dissociable cognitive strategies for sensorimotor learning. Nat Commun 10:40.

Morehead JR, Qasim SE, Crossley MJ, Ivry R (2015) Savings upon Re-Aiming in Visuomotor Adaptation. J Neurosci 35:14386–14396.

Orban de Xivry JJ, Lefevre P (2015) Formation of model-free motor memories during motor adaptation depends on perturbation schedule. J Neurophysiol 113:2733–2741.

Paz R, Boraud T, Natan C, Bergman H, Vaadia E (2003) Preparatory activity in motor cortex reflects learning of local visuomotor skills. Nat Neurosci 6:882–890.

Pellizzer G, Georgopoulos AP (1993) Common processing constraints for visuomotor and visual mental rotations. Exp Brain Res 93:165–172.

Perich MG, Gallego JA, Miller LE (2018) A Neural Population Mechanism for Rapid Learning. Neuron 100:964–976 e967.

Pidoux L, Le Blanc P, Levenes C, Leblois A (2018) A subcortical circuit linking the cerebellum to the basal ganglia engaged in vocal learning. Elife 7:e32167.

Rueda-Orozco PE, Robbe D (2015) The striatum multiplexes contextual and kinematic information to constrain motor habits execution. Nat Neurosci 18:453–460.

Sack AT, Lindner M, Linden DEJ (2007) Object- and direction-specific interference between manual and mental rotation. Perception & Psychophysics 69:1435–1449.

Savoie FA, Thenault F, Whittingstall K, Bernier PM (2018) Visuomotor Prediction Errors Modulate EEG Activity Over Parietal Cortex. Sci Rep 8:12513.

Schouten JF, Bekker JA (1967) Reaction time and accuracy. Acta Psychol (Amst) 27:143–153.

Schween R, Taube W, Gollhofer A, Leukel C (2014) Online and post-trial feedback differentially affect implicit adaptation to a visuomotor rotation. Exp Brain Res 232:3007–3013.

Seidler RD, Noll DC, Chintalapati P (2006) Bilateral basal ganglia activation associated with sensorimotor adaptation. Experimental Brain Research 175:544–555.

Shadmehr R, Mussa-Ivaldi FA (1994) Adaptive representation of dynamics during learning of a motor task. J Neurosci 14:3208–3224.

Shadmehr R, Holcomb HH (1997) Neural correlates of motor memory consolidation. Science 277:821–825.

Shadmehr R, Holcomb HH (1999) Inhibitory control of competing motor memories. Experimental Brain Research 126:235–251.

Tanaka H, Sejnowski TJ, Krakauer JW (2009) Adaptation to visuomotor rotation through interaction between posterior parietal and motor cortical areas. Journal of Neurophysiology 102:2921–2932.

Taylor JA, Thoroughman KA (2007) Divided attention impairs human motor adaptation but not feedback control. J Neurophysiol 98:317–326.

Taylor JA, Thoroughman KA (2008) Motor adaptation scaled by the difficulty of a secondary cognitive task. PLoS One 3:e2485.

Taylor JA, Ivry RB (2011) Flexible cognitive strategies during motor learning. PLoS Comput Biol 7:e1001096.

Taylor JA, Ivry RB (2014) Cerebellar and prefrontal cortex contributions to adaptation, strategies, and reinforcement learning. In: Progress in Brain Research, pp 217–253: Elsevier.

Toni I, Ramnani N, Josephs O, Ashburner J, Passingham RE (2001) Learning arbitrary visuomotor associations: temporal dynamic of brain activity. Neuroimage 14:1048–1057.

von Helmholtz H, Southall JP (1924) Helmholtz’s treatise on physiological optics, Vol. 1, Trans.

Yin HH, Knowlton BJ (2006) The role of the basal ganglia in habit formation. Nat Rev Neurosci 7:464–476.

Zach N, Kanarek N, Inbar D, Grinvald Y, Milestein T, Vaadia E (2005) Segregation between acquisition and long-term memory in sensorimotor learning. European Journal of Neuroscience 22:2357–2362.

Zarahn E, Weston GD, Liang J, Mazzoni P, Krakauer JW (2008) Explaining savings for visuomotor adaptation: linear time-invariant state-space models are not sufficient. J Neurophysiol 100:2537–2548.

Zhou X, Tien RN, Ravikumar S, Chase SM (2019) Distinct types of neural reorganization during long-term learning. J Neurophysiol 121:1329–1341.

